# The DMCdrive: practical 3D-printable micro-drive system for reliable chronic multi-tetrode recording and optogenetic application in freely behaving rodents

**DOI:** 10.1101/2020.04.24.059394

**Authors:** Hoseok Kim, Hans Sperup Brünner, Marie Carlén

**Affiliations:** Department of Neuroscience, Karolinska Institutet, 171 65 Stockholm, Sweden; Department of Biosciences and Nutrition, Karolinska Institutet, 141 83 Huddinge, Sweden

## Abstract

Electrophysiological recording and optogenetic control of neuronal activity in behaving animals have been integral to the elucidation of how neurons and circuits modulate network activity in the encoding and causation of behavior. However, most current electrophysiological methods require substantial economical investments and prior expertise. Further, the inclusion of optogenetics with electrophysiological recordings in freely moving animals remains a general challenge. Expansion of the technological repertoire across laboratories, research institutes, and countries, demands open access to high-quality devices that can be built with little or no prior expertise from easily accessible parts of low cost. We here present a very affordable, truly easy-to-assemble micro-drive for electrophysiology in combination with optogenetics in freely moving mice and rats. The DMCdrive is particularly suited for reliable long-term recordings of neurons and network activities, and simplify optical tagging and manipulation of neurons in the recorded brain region. The highly functional and practical drive design has been optimized for accurate tetrode movement in brain tissue, and remarkably reduced build time. We provide a complete overview of the drive design, its assembly and use, and proof-of-principle demonstration of long-term recordings paired with cell-type-specific optogenetic manipulations in the prefrontal cortex (PFC) of freely moving transgenic mice and rats.

## Introduction

Tetrode-containing micro-drives have long been an important technique for recordings of extracellular neuronal signals in behaving mice^1^ and rats^2^. More recently developed techniques, such as neuronal population calcium imaging^3^ and high-density silicon probes^4^ are superior to micro-drive arrays in respect to the number of neurons that can be recorded simultaneously. Despite this, there is still a range of application areas where micro-drive arrays are the method of choice, particularly for chronic recordings of action potential and local field potential (LFP) activity in freely moving rodents, including in conjunction with optogenetics. We here present a low-cost (~10 $/drive), easy to assemble microdrive for long-term stable tetrode recordings in conjunction with optogenetics (the DMCdrive). The strategy for the design has been to construct a user-friendly low size-low weight drive that reliably provides a) single-unit recordings over weeks, b) persistent precise and predictable movement of all tetrodes c) efficient opto-tagging and optical manipulation of neurons. The drive consists of a few lab-made parts, printed using a standard 3D printer, a custom-made electronic interface board (EIB), and parts available off-the-shelf, and the easy-to-assemble design makes the drive particularly suitable for researchers with no or little experience of drive building. The drive is designed for movement (in depth) of the tetrodes at any time, an approach holding several advantages. First, as the neuronal signal deteriorates over time (likely due to the formation of a glial sheath that shields the electrodes from active neurons^5,6^), moving of the electrodes will in many cases restore the signal and provide stability of the recording conditions^6,7^. Second, adjustment of the depth of recording electrodes enables the recording of a new set of neurons, increasing the total number of neurons recorded per animal. However, if the moving force fails to move the electrodes through the glial sheath, the neural signal cannot be restored^8^, and new neurons cannot be recorded. For this reason, reliable movement of all tetrodes was of high priority in the design of the drive, and by moving all tetrodes together our drive maximizes the moving force. Further, for precise targeting of recordings to specific locations, and estimation of the recording location after moving of the tetrodes, accurate depth movement of the tetrodes is of high importance. To achieve this, the movement of the tetrodes is controlled by a single, easily accessible shuttle screw, reliably moving the tetrodes 350 μm/360°. Optogenetics in conjunction with extracellular recordings is a powerful method to study neural circuit function^10^. Optical tagging of neurons expressing light-activated opsins enables identification of the activity of select neurons, e.g., of specific neuron types^11,12^ or specific projections^13^. In addition, the inclusion of optogenetics allows for the manipulation of a neuronal population and recording of the effects on single-unit and/or LFP activities in the local network^14^. However, despite great technical advances it remains a challenge to combine optogenetics with extracellular electrophysiology in freely behaving animals. The DMCdrive integrates long-term recordings with highly reliable manipulation of light-sensitive neurons. We provide a complete overview of the design of the drive, its assembly and use. For a demonstration of the utility of the drive, data from long-term recordings in the PFC of freely moving transgenic mice and rats is provided, and includes opto-tagging of prefrontal inhibitory interneurons expressing parvalbumin (PV) and optogenetic entrainment of prefrontal gamma oscillations.

## Results

### Fabrication and assembly of the DMCdrive

The DMCdrive consists of 3D-printed parts, commercially available parts, and a custom-made EIB designed to support 16 channels/4 tetrodes and one optical fiber (**Fig. 1A, B**). All required design files and a list of the materials, small parts, and tools needed for building of the drive are freely available at http://carlenlab.org/data. The focus of the drive design has been on functional fidelity, but also on very fast assembly, and minimization of the size and weight (16 x 16 x 18 mm, width x length x height; weight < 2 g; **Fig. 1C**). D-printing, drive assembly, tetrode and fiber optic loading, and testing of the impedance of the tetrodes can be achieved in 4 hours with some practice.

**Figure 1.**
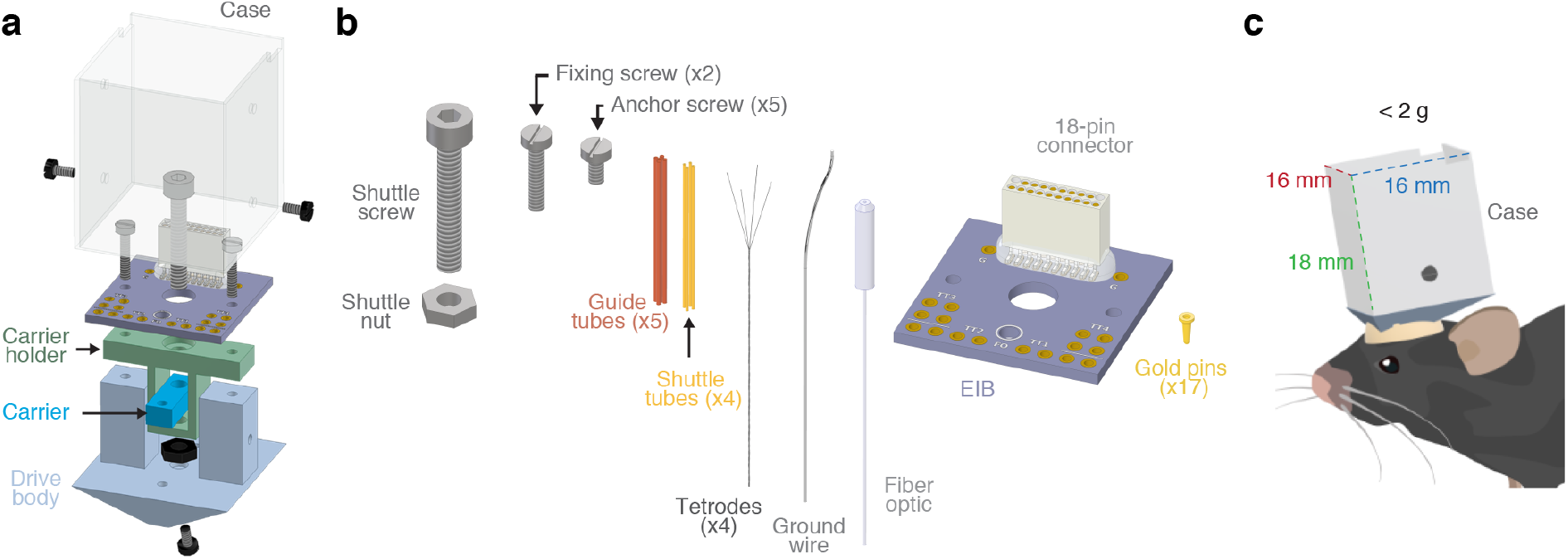
Schematic of the DMCdrive and component parts. **A**) Isometric illustration of the functional parts of the DMCdrive. Four of the six 3D-printed parts are indicated (case, carrier holder, carrier, and drive body, respectively). **B**) Parts needed, in addition to the 3D-printed parts, to build a DMCdrive for multi-tetrode recordings in conjunction with optogenetic manipulations. **C**) Illustration of a mouse implanted with a DMCdrive. Outside measurements of the case: height; 18 mm (green dotted line), width and length; 16 mm (red and blue dotted line, respectively). Assembled with four tetrodes, an EIB, a ground wire and an optical fiber the drive weighs approximately 2 g.

The six 3D-printed parts, designed using a 3D CAD modeling software (Autodesk Inventor, Autodesk), can be printed using a standard, low-cost 3D printer in less than 2 hours (**Fig. 2A**). The 3D-printed parts are printed together in one structure and should be cleaned from residual debris and structure (e.g., using an art knife). Screw holes in the printed drive parts should be carefully cleaned by drilling with appropriately sized drill bits (**Fig. 2B**).

**Figure 2.**
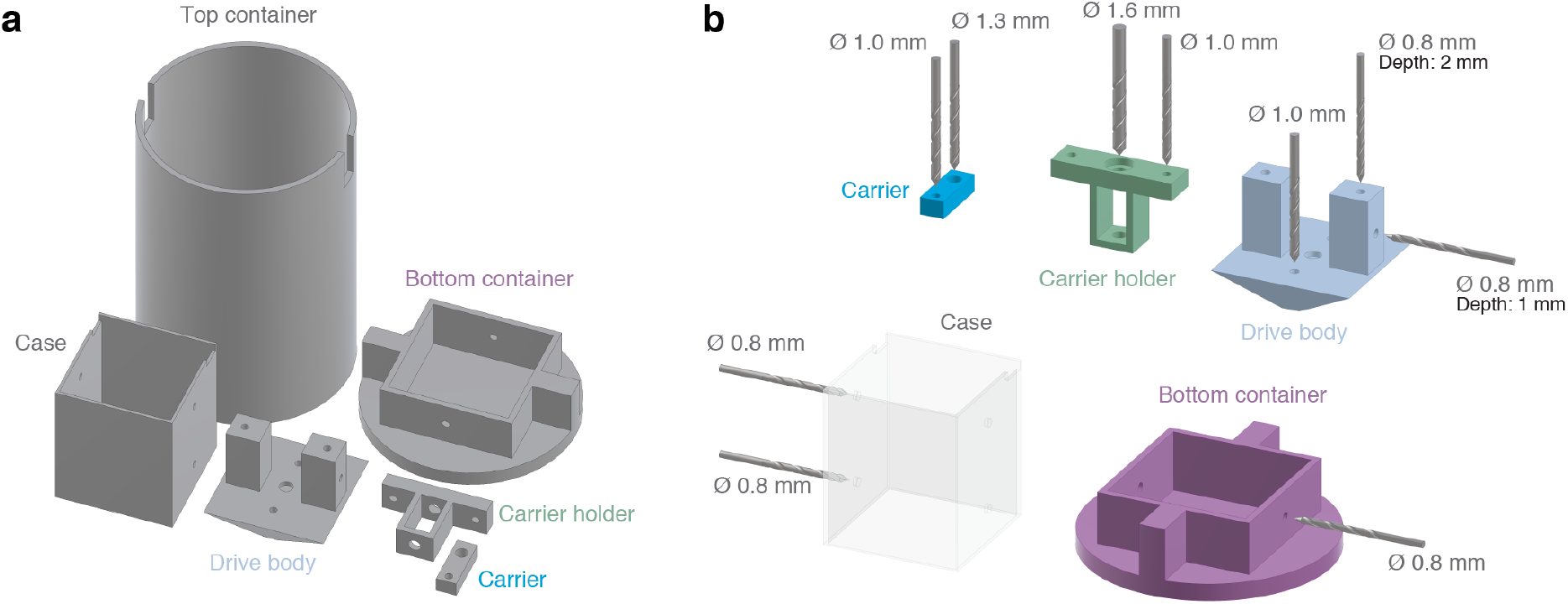
3D printing. **A**) All 3D-printed parts of the DMCdrive, in a layout suitable for printing. The parts can be printed in less than two hours using a normal desktop 3D printer (Makerbot Replicator, Makerbot). **B**) To ensure the correct sizing of screw holes, drilling of the holes is recommended after printing.

The drive assembling starts by building of the carrier assembly, which includes securing of the tetrode/ fiber-carrier to the carrier-holder using a shuttle screw and a screw nut (**Fig. 3A, B**). The carrier assembly is thereafter placed in the drive body (generating the drive assembly), and the EIB holding an 18-pin Omnetics connector is ‘subsequently connected to the drive assembly with two fixing screws (**Fig. 3C, D**). An anchoring screw added to the bottom of the drive (**Fig. 3D**) is used for stable fixation of the drive to the dental cement surrounding the craniotomy.

**Figure 3.**
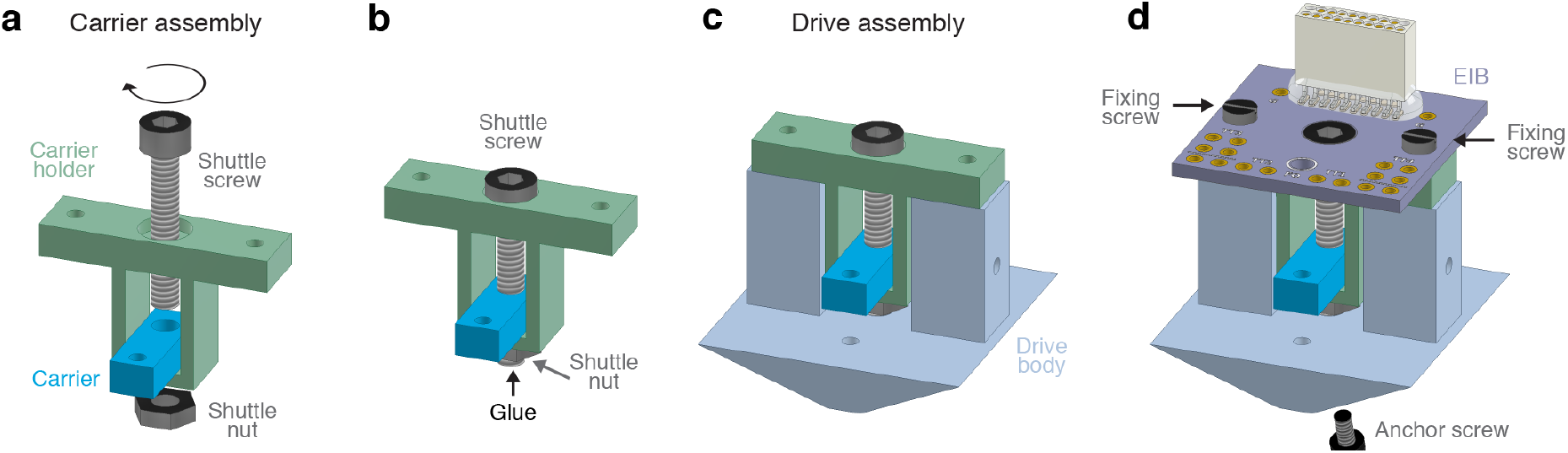
Drive assembly. **A**) Insert the shuttle screw (M1.6 x 10 mm, hex socket head) into the carrier holder to connect the carrier with the carrier holder (‘carrier assembly’). Install the shuttle nut (M1.6, hex nut) on the shuttle screw. **B**) Secure the shuttle nut to the shuttle screw with glue (black arrow). **C)** Place the carrier assembly in the drive body (‘drive assembly’). **D)** Connect an EIB holding an 18-pin Omnetics connector to the drive assembly with two fixing screws (M1 x 5 mm thread length). Add an anchor screw (M1 x 2 mm thread length) to the bottom of the drive body for later securing of the drive to the dental cement surrounding the craniotomy.

For the use of four tetrodes and one optical fiber, a pre-assembled bundle of five guide tubes is inserted through, and secured to, the drive body, and the carrier (**Fig. 4A, B).** After cutting of the guide tubes into two parts (**Fig 4C**), shuttle tubes are inserted into four of the guide tubes, and thereafter secured (**Fig. 4D**). One or two ground wires are inserted through the drive body and connected to the EIB with gold pins (**Fig. 4E, F**). The pre-fabricated tetrodes^15^ are carefully loaded into the shuttle tubes one by one, connected to the EIB using gold pins, and glued to the top of the shuttle tubes (**Fig. 4E, F**). For the inclusion of optogenetics, an optical fiber coupled to a fiber ferrule^16^ is inserted through the fifth guide tube (**Fig. 4G**) and firmly attached to the EIB (**Fig. 4H).** The tetrode bundle is then cut to the desired length (**Fig. 4H**). For the protection of the drive, the 3D printed case is placed over the assembled drive and secured using two anchor screws (**Fig. 4I**). As a final step the tips of the tetrodes are gold-plated to lower their impedance^15^. The finished drive can be placed in the 3D-printed container for safe storage and transportation (**Fig. 4J**).

**Figure 4.**
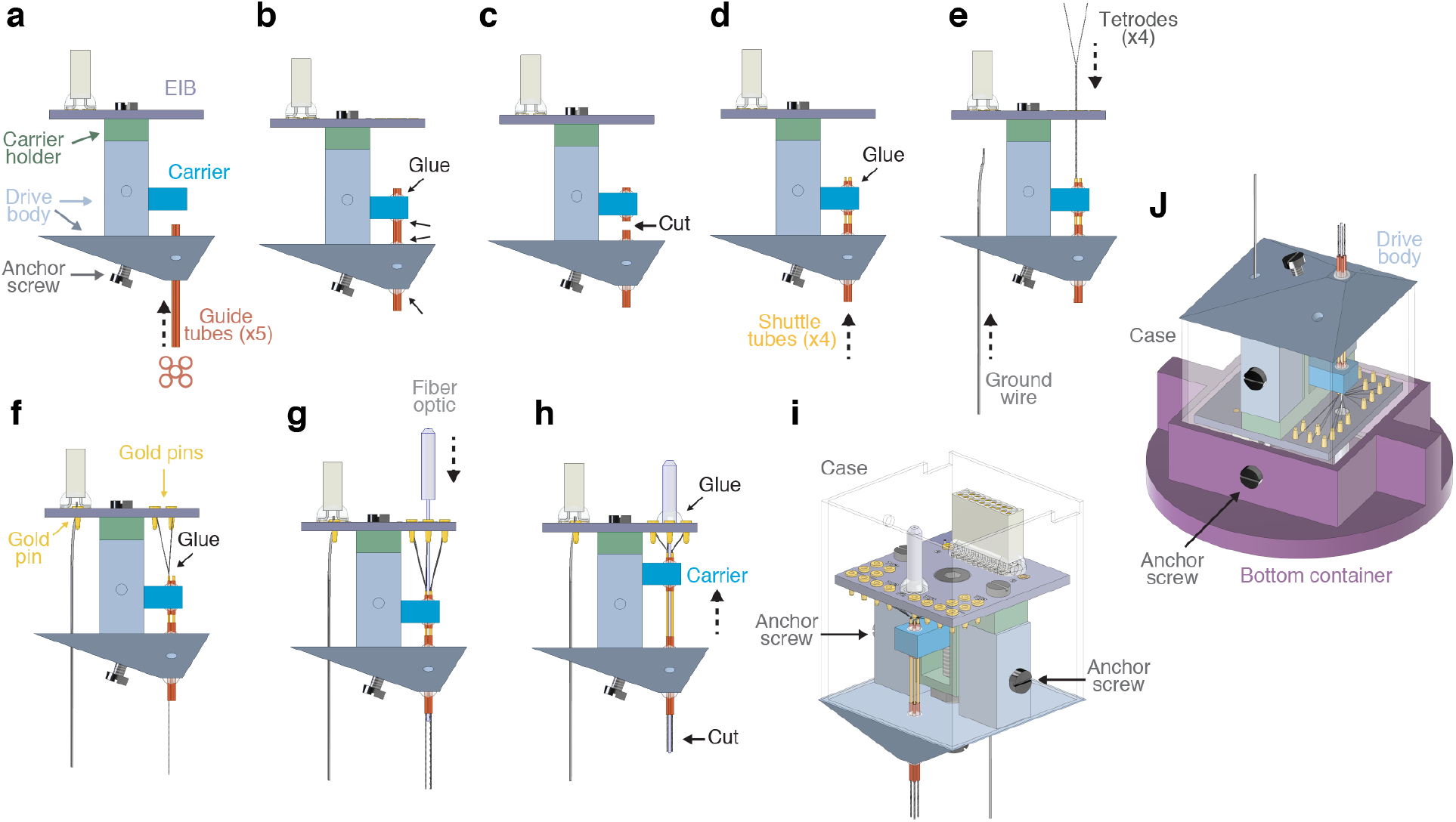
Tetrode and fiber optic loading. **A**) Side view of the DMCdrive. Start by moving the carrier to the bottom of the drive. Insert the five guide tubes (0.0120”/ 0.0140”, ID/OD) into the drive body. The configuration of the guide tubes is dictated by the desired arrangement of the tetrodes and the optical fiber (e.g., round or flat). **B**) Glue (arrows) the guide tubes to the drive body and carrier. **C**) Cut (arrow) the guide tubes between the drive body and the carrier. **D**) Insert four shuttle tubes (0.0049”/0.0064”, ID/OD) into four of the guide tubes. The fifth guide tube will hold the optical fiber. Glue (arrow) the shuttle tubes to its respective guide tube. **E**) Insert the ground wire, without insulation at the ends, into and through the drive body from below. Place one tetrode in each shuttle tube. **F**) Secure the ground wire and the tetrodes to the EIB using gold pins. To enable movement of the tetrodes, glue (arrow) the tetrodes to its respective shuttle tube. **G**) Insert the fiber optic (fiber: multimode, 200 μm core, 0.22-0.5 NA; ferrule: Ø1.25 mm, 6.4 mm long, Ø230 μm bore; Thorlabs) through the EIB into the empty guide tube. **H**) Glue (arrow) the fiber optic to the EIB. Move the carrier up as far as possible. **I**) Place the assembled drive in the case, and secure the drive to the case using two anchor screws (‘finished drive’). **J**) For storage and transport of the drive, place the finished drive upside down in the bottom container and secure with anchor screws, and then place the top container over the drive.

### Chronic neuronal recordings in mice and rats

To evaluate the performance of the DMCdrive for neurophysiological recordings in conjunction with optogenetics in freely behaving animals, adeno-associated viral (AAV) vectors with Cre-dependent expression of channelrhodopsin-2 (ChR2) were targeted to the left (1 mouse and 2 rats) or right (1 mouse and 1 rat) medial PFC (mPFC) of transgenic PV-Cre mice and PV-Cre rats (**Methods**). 2-3 weeks later the animals were implanted with a DMCdrive at the same location, with the tetrodes targeted to the prelimbic cortex (**Fig. 5A, B**). After 7 days of recovery from surgery, the animals were placed in an open field area (25×25 cm) and the activity of mPFC neurons was recorded (5-30 min).

**Figure 5.**
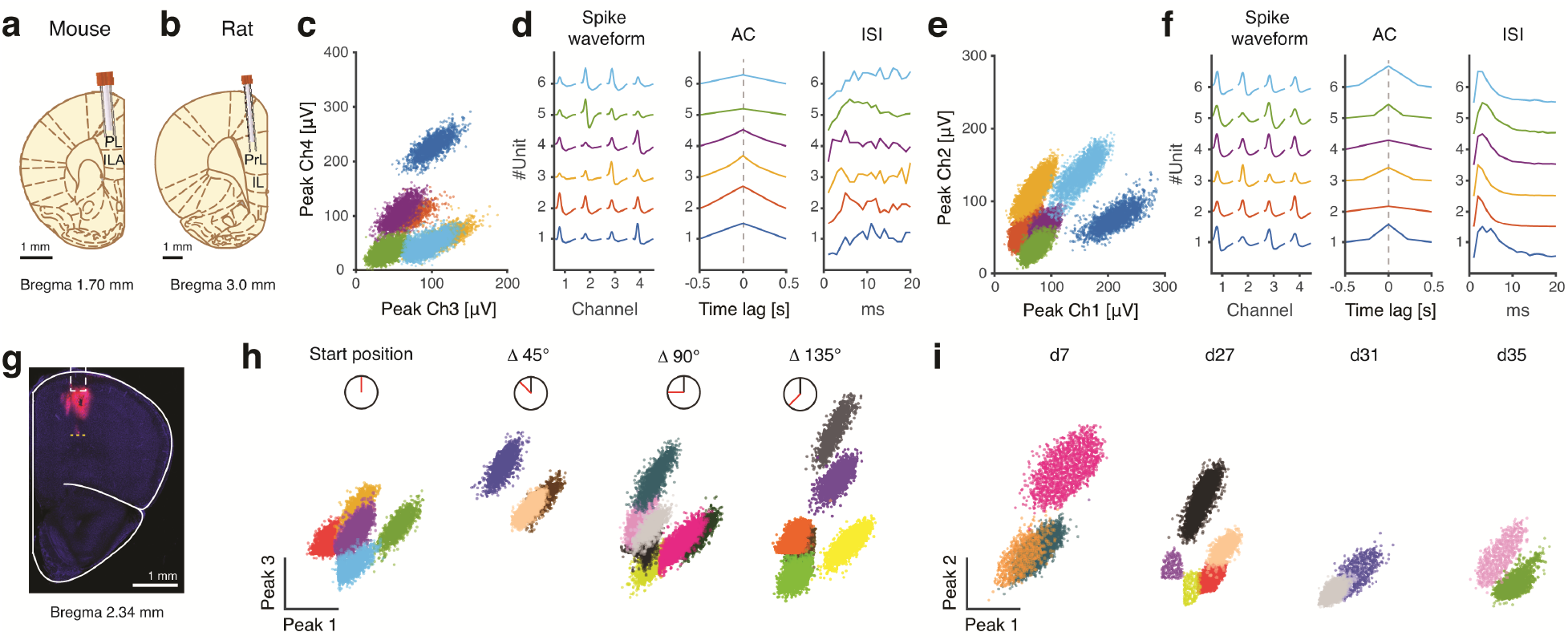
Chronic extracellular single-unit recordings in awake animals. **A-B**) Schematic illustration of a coronal section with tetrodes and an optical fiber targeted to the mPFC in a PV-Cre mouse and PV-Cre rat, respectively. **C**) Peak amplitude plot of action potentials recorded from channel 3 and 4 of a tetrode implanted in a mouse, with colorcoding of the identified clusters. **D**) Average spike waveforms of the single-unit clusters in (C), as recorded on the four electrodes of the tetrode. Autocorrelation (AC) and inter-spike interval (ISI) confirm that the spike times of each cluster belong to individual neurons. **E**) Peak amplitude plot of action potentials recorded from channel 1 and 2 of a tetrode implanted in a rat, with color-coding of the identified clusters. F) Average spike waveforms of the single-unit clusters in (E), as recorded on the four electrodes of the tetrode. AC and ISI confirm that the spike times of each cluster belong to individual neurons. **G**) Tetrode mapping with CM-Dil (red) and DAPI (blue) in the mouse mPFC. Dashed white box: location of implanted tetrodes and an optical fiber. Yellow dashed line: the ventral limitation of CM-Dil labelling, corresponding to the end location of the tetrode tips after tetrode lowering by three full turns of the shuttle screw. **H-I**) Single-unit recordings in the mPFC of freely moving rats. **H**) Example recordings from a tetrode that was lowered in steps (5 minutes apart) by 45° turns of the shuttle screw (45° turn: ~44 mm). At each depth (n = 4) neuronal signals were recorded, and spikes were clustered (5, 3, 7, and 5 well-isolated single units, respectively) and plotted by the peak amplitude as recorded on electrode 1 and 3 of the same tetrode. **I)** Example recordings from a tetrode discriminating single units 35 days post-implantation. Recordings were performed at 19 different days. The tetrodes were lowered by a 45° turn of the shuttle screw after each recording session. Peak

Analysis of spiking data from single tetrodes in both mice and rats revealed well-separated clusters (**Fig. 5B-F**), and recording of up to 14 single units from all tetrodes in a single recording session (data not shown). Precise placement and stable control of the tetrodes were important priorities in the design of the drive, as was the possibility to record new neurons over time. To illustrate the reliability of the tetrode movement in awake animals, we performed an *in vivo* experiment in which the tips of the tetrodes and the optical fiber were dipped in a red fluorescent dye (CM-Dil) before implantation, for visualization of the tetrode track (n = 2 mice). Using the optical fiber tip as a reference point, the shuttle screw was turned one full turn three times (with appropriate time intervals between every turn), corresponding to the movement of the tetrodes 3 x 350 μm = 1050 μm (**Fig. 5G**). Processing of the tissue revealed DiI labelling along the tetrode track all the way down to the final tetrode position.

To confirm that moving of the tetrodes results in reliable detection of new units, four short (5 minutes) successive recordings sessions were performed over the course of 50 minutes (n = 1 rat). After every recording the shuttle screw was turned counter-clockwise 1/8 of a turn (~45°), corresponding to the lowering of the tetrodes ~44 μm. With the use of a manipulator (hex screwdriver) the tetrodes can be moved vertically without risk of damaging of the tetrodes. Further, compared to existing micro-drives holding multiple independently movable tetrodes^7,17^, our single screw-based drive system limits the time used for tetrode movement, reducing the stress put on the animal. Every turn of the screw rendered a distinct electrophysiological condition around the tetrode tip comparable to the conditions of the previous recording (before the turn), with a different number and characteristics of the spiking in each recording, strongly indicative of new neurons being recorded in every session (**Fig. 5H**). In a different experiment, recordings (followed by lowering of the tetrodes) were performed in up to 19 different days, with a successful recording of single units every day, including 35 days after implantation (the longest time point tested; **Fig. 5I**, n = 3 rats). Together this set of experiments confirm efficient control of the tetrodes and long-term recordings using the DMCdrive.

### Optogenetic modulation of single-unit and LFP activity

Cortical PV interneurons are important circuit regulators, and experiments employing optogenetics in intact mice have been central to demonstrate the role of this neuronal population in the generation of gamma oscillations, circuit functions, cognition and behavior^14,18–20^. Here we combined recordings with opto-tagging and optogenetic drive of PV interneurons in the mPFC in mice and rats using the DMCdrive. Application of 40 Hz blue light (473 nm, 3 ms light pulse, 1 s duration, 5-10 mW) resulted in a significantly increased short-latency firing in a set of neurons in both rats and mice (**Fig. 6A-C**). The light-evoked action potentials were characteristic of prefrontal fast-spiking (FS) PV interneurons, with short half-trough width and comparable peak and trough deflections^18^ (**Fig. 6D**). Comparison of spontaneous and light-activated action potentials revealed very high similarity (99.7%) between the spike waveforms^21^, demonstrating true light-activation of ChR2-expressing PV interneurons (**Fig. 6D**). Prefrontal inhibitory PV interneurons provide strong inhibition onto excitatory neurons in the local network^18^, and in agreement with this application of blue light resulted in silencing of nearby neurons with waveform properties typical of cortical excitatory neurons^14,22^ in both mice and rats (**Fig. 6E-H**).

**Figure 6.**
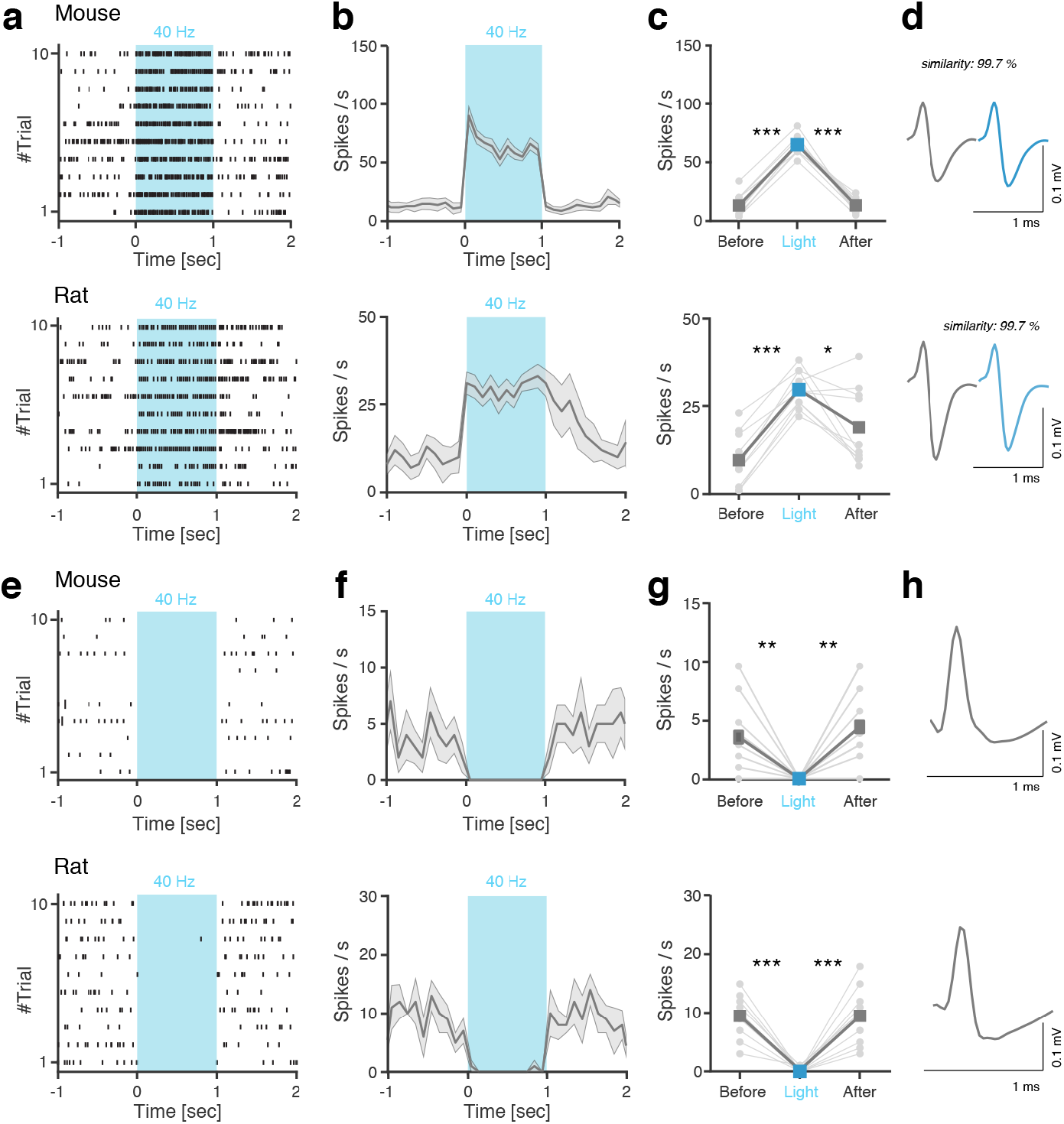
Optogenetics in conjunction with chronic single-unit recordings. (**A-H)** Optogenetic manipulation of ChR2-expressing mPFC PV interneurons in PV-Cre mice and PV-Cre rats. Top: mouse, bottom: rat. **A-D**) Light-activated mPFC PV interneurons. **A**) Spike raster of an example light-responsive PV interneuron responding to 40 Hz blue light (473 nm, 1 s, 3 ms pulses) application with significantly increased spiking. **B-C**) Average modulation of the neuron across 10 trials. **D**) Spike waveform of the recorded neuron. Spontaneous and light-evoked action potential waveforms exhibit very high similarity. **E-H**) Concurrently recorded WS putative principal neurons in the local mPFC circuitry. **E**) Spike raster of an example WS putative principal neuron demonstrating significantly decreased spiking in response to light-activation of local PV interneurons. **F-G**) Average modulation of the neuron across 10 trials. **H)** Spike waveform of the recorded neuron. Shaded area and error bars: ± SEM. *p < 0.05, **p < 0.01, ***p < 0.001 by paired t-test.

The LFP activity provides information about the activity of aggregates of neurons, including the temporal scales of the activities^23^. It has been firmly established that synchronous activity of cortical PV interneurons generates gamma oscillations (30-80 Hz) in the local network^24,25^. Optogenetic studies have shown that activation of PV interneurons results in an increase of the LFP power specifically at gamma frequencies^14^. We here activated prefrontal PV interneurons with blue light at a typical gamma frequency (40 Hz, 3 ms light pulse, 1 s duration, 5-10 mW) in freely moving mice and rats. Analysis of the LFP revealed increased LFP power and gamma activity time-locked to the light application (**Fig. 7A, B**). In agreement with the mechanisms of fast-spiking PV interneurons generating cortical gamma rhythms, light-activated neurons with increased firing displayed typical narrow-spiking (NS) features (**Fig. 6D**).

**Figure 7.**
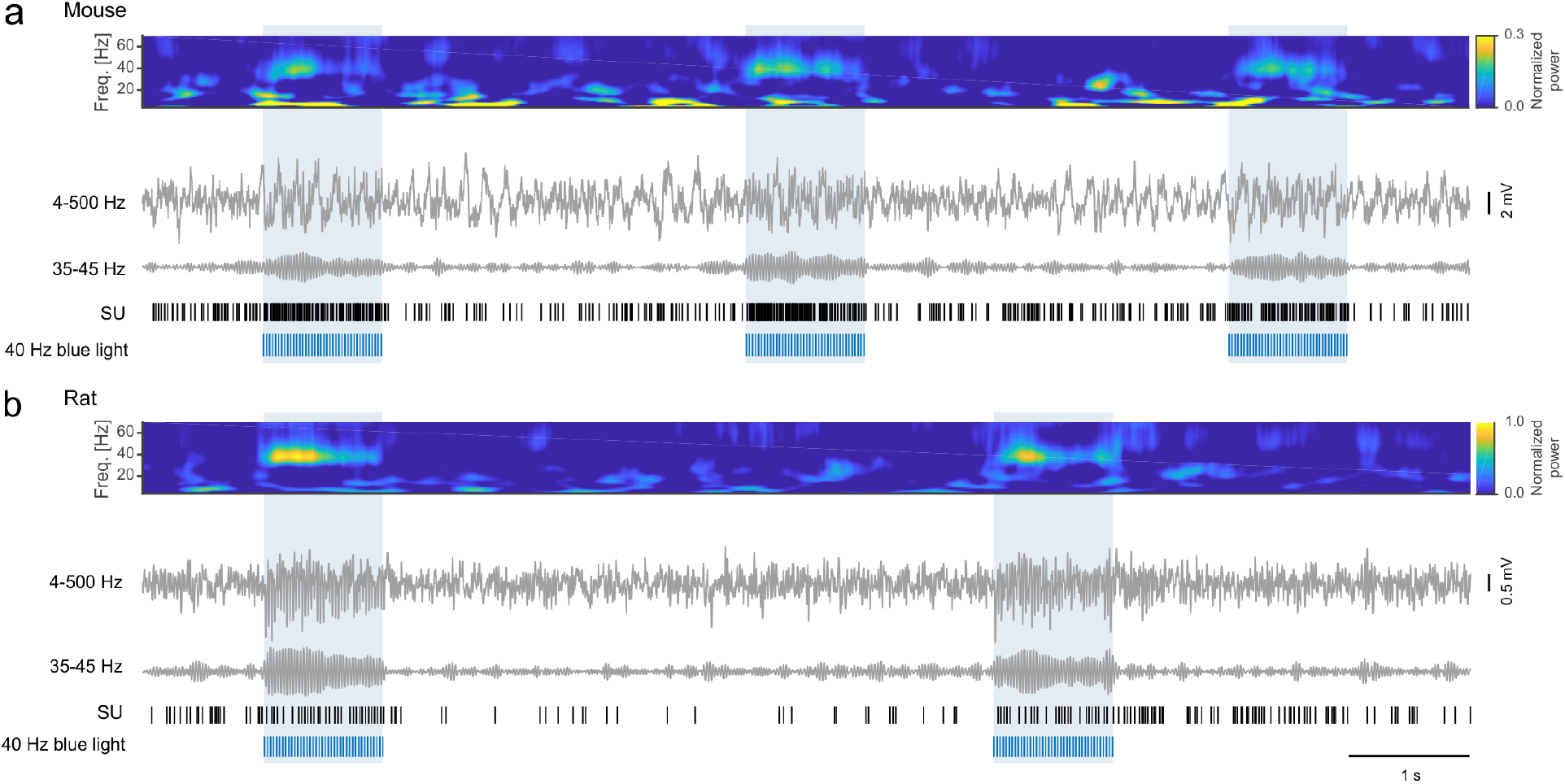
Optogenetics in conjunction with LFP recordings. **A-B**) Optogenetic manipulation of ChR2-expressing mPFC PV interneurons. Spectrogram (4-70 Hz), raw LFP (4-500 Hz), band-pass filtered LFP (35-45 Hz), and single-unit (SU) activity from mPFC recordings (10 s) in freely moving PV-Cre mice **(A)** and PV-Cre rats (**B**). Blue light application (473 nm, 1 s, 40 Hz, 3 ms pulses) results in increased oscillatory activity in the gamma range as seen in the LFP. Same SU as in Figure 6A.

## Discussion

Electrophysiological recordings in combination with local optogenetic manipulations in freely behaving animals allow investigation of neuronal mechanisms underlying brain functions, and have been used to probe emotion, memory, attention, and perception, and more^18,20,26,27^. The ability to tag neurons with light for identification of activity patterns of select neuronal populations has greatly aided the understanding of how specific neuron types contribute to behavior ^28,29^. Tetrode arrays have long been the gold standard for *in vivo* neuronal recordings and provide high temporal (sub-millisecond) and spatial (single-unit) resolution. Micro-drive systems with movable tetrodes enable high-yield and long-term recordings over the course of days to weeks, and the integration of fiber optics allows control of neurons or neural circuits during recordings. Application in freely behaving rodents puts constraints on the size and weight on the implant, and as a general rule an implant can only be used in a single animal. Implants, thus, must regularly be fabricated or purchased.

We here present an affordable simple-to-operate, truly easy-to-build, compact and low-weighted micro-drive for reliable multi-tetrode electrophysiological recordings in combination with optogenetic investigations. The DMCdrive consists primarily of 3D-printed components, remarkably reducing the cost of the device. The 3D-printing in addition provides possibilities for desired customizations. While alternative implant designs report low fabrication costs^30,31^, the ~10USD/ DMCdrive includes not only six 3D-printed parts but also an EIB with an 18-pin Omnetics connector, eight screws and a nut, making our drive design a genuinely affordable alternative for many laboratories. The 3D-printed drive parts are configured for very straightforward assembly and reduced drive build time. For reduced friction between the tetrodes and the guide tubes, and predictable vertical linear motion of the tetrodes, all tetrodes are disposed parallel to each other and driven simultaneously by a large core screw (hex socket screw) in our drive design. Further, there is no physical contact between the tetrodes and the drive manipulator (hex screwdriver) used to move the tetrodes. Our design does not support individually movable tetrodes, which is often the method of choice for high-yield recording of neurons from multiple brain structures and layers, but holds the advantage of highly precise targeting of brain regions without tetrode bending and with limited tissue damage. Thus, our drive system can provide superior stability of the recording conditions over time.

The presented design is optimized for chronic recordings in freely moving mice and includes only four tetrodes, but the drive design can flexibly be modified to accommodate up to eight tetrodes suitable for multi-site recordings, or larger scale recordings in rats. As demonstrated, the original design can nevertheless be proficiently used in rats, although rats would be able to carry larger designs holding more tetrodes. The size and weight of the drive have been reduced to a minimum, limiting the effects on animals’ behavior, and the rectangular shape of the implant is particularly convenient for lowering of tetrodes and connecting/disconnecting cables in awake animals. The main possible limitation of the drive design is the permanent fixation of the fiber optic to the device, prohibiting any movement of the optical fiber. As the choice of opsin gene, the functional expression levels of the opsin, and the light power density reaching the light-sensitive neurons all are factors influencing the efficacy of light-based control of neuronal activity, the experimental parameters in individual experiments will determine how far from the optical fiber the tetrode tips can be lowered with preserved optical manipulations^32^. Integration of movable optical fibers into the recording array, while avoiding increases in the size, weight, and complexity of the drive, yet imposes a great technical challenge for *in vivo* electrophysiologists. We here prioritized other important parameters in the design of the drive.

To date, various off-the-shelf and in-house-built devices to support chronic multiple tetrode recordings together with optical stimulation have been designed, including the VersaDrive (Neuralynx), systemDrive, SLIQ hyperdrive, flexDrive, and more^7,33,34^. Compared to the DMCdrive, these devices can carry a larger number of recording channels and allow positioning of each tetrode independently. However, as the number of tetrodes increases, so does the size and weight of the implant, which needs to be accounted for in the planning of experiments in freely moving animals. Further, the complexity of a drive greatly affects the time and effort needed for tetrode/fiber optic loading and building of the device. In addition, the bending of the tetrodes and the concentric arrangement of tetrode bundles in many current devices are more likely to impose friction between a tetrode and the guide tube during tetrode lowering, negatively impinging on the reliability of tetrode movement.

Given the outlined, and highly intentional, advantages the DMCdrive offers, we firmly believe that the outlined drive is a welcomed addition to the toolbox in experimental neuroscience and will find many application areas, particularly for users with financial constraints or limited prior experience of neurophysiology in behaving rodents.

## Methods

### Animals and surgical procedures

All procedures were approved and performed in accordance and compliance with the guidelines of the Stockholm Municipal Committee for animal experiments. Adult PV-Cre mice (males, 3-4 months old; Jax Stock no. 008069; https://jax.org/strain/008069) and PV-Cre rats (manuscript in preparation; male and female, 2-4 months old) were used. Animals were housed under a standard 12-hour light/dark cycle. For viral injections, the animals were injected with buprenorphine (0.05 mg/kg s.c.) 30 min prior to surgery. Animals were anesthetized with ~2% Isoflurane in oxygen, fixed in a stereotaxic device, and lidocaine (maximum 5mg/kg) was injected locally before skin incision. A small craniotomy was unilaterally made above the mPFC (mice: 1.7 mm anterior to Bregma and 0.25 mm lateral to midline, rats: 3.0 mm anterior and 0.5 mm lateral to the midline). An adeno-associated viral vector with Cre-dependent expression of ChR2, AAV-DIO-ChR2-mCherry (rAAV5/EF1a-DIO-hChR2(H143R)-mcherry; 4×10e12 viral particles/ml, UNC Vector Core), was used for optogenetic activation of PV-expressing interneurons. The virus (0.5 μl) was injected into the mPFC, centered to the prelimbic area, (mice: AP 1.7 mm, ML 0.25 mm, DV −1.0 mm; rats: AP 3.0 mm, ML 0.5mm, DV −2.0 mm) using a glass micropipette coupled to a motorized stereotaxic injector (Stoelting) at a rate of 0.1 μl/min. The glass pipette was kept in place for 5 min after injection and then slowly withdrawn to minimize backflow. The skin was closed with tissue glue (Vetbond, 3M) or sutures (silk, Ethicon). Carprofen (5 mg/kg) was administered subcutaneously to the animals before recovery from anesthesia, and 24h after surgery. Three weeks after virus injection, the animals were implanted with a DMCdrive (identical analgesia and anesthesia protocols as for virus injection). The implant was positioned in the craniotomy made prior to the virus injection. The craniotomy was covered with vaseline to protect the tetrodes and the optical fiber. Stainless steel skull screws (3 for mice and 5 for rats) and dental cement were used for firm attachment of the drive to the skull. The ground wire (uncovered part) of the drive was wrapped around the skull screws as ground. To protect the EIB and the exposed parts of the drive from physical contact with the environment, the top of the implant (the case) was sealed with clear office tape.

### Determining tetrode placement

To illustrate the intended control and precision of depth movement of the tetrodes in freely moving animals, a DMCdrive was implanted in the dorsal prefrontal cortex of a mouse. Prior to the implantation, the tetrodes tips were aligned with the optical fiber, and carefully dipped in CellTracker^TM^ CM-Dil fluorescent dye (ThermoFisher). 24 hours post-surgery, the tetrodes were moved down ~1 mm by three full (360°) turns of the shuttle screw (with appropriate time intervals between every turn, corresponding to movement of the tetrodes 3 x 350 μm = 1050 μm). One hour later the mouse was transcardially perfused with 4% PFA in PBS. 300 μm thick brain sections were cut on a vibratome (Leica VT1000, Leica Microsystems). Brain sections were cleared according to a modified CUBIC clearing method ^35^. Z-stack images were acquired at 10x, using a Zeiss (LSM800) confocal laser scanning microscope.

### Neural data acquisition and optogenetic manipulation

Single-unit and LFP activities were recorded using a Cheetah data acquisition system (Digital Lynx 4SX, Neuralynx). Unit signals were amplified (x10,000), band-pass filtered (600-6,000 Hz), digitized (32 kHz), and stored on a PC for off-line analysis. LFP signals were acquired from one channel of each tetrode at 32 kHz sampling frequency, and band-pass filtered between 0.1 and 500 Hz. For optogenetic experiments, blue light (473 nm, 3 ms pulse width, 1 sec duration, 5-10 mW at the fiber tip, 40 Hz) was delivered through the optical fiber using a DPSS laser (Cobolt).

### Spike sorting

To identify single units from tetrode data, spikes were manually sorted based on various spike waveform features (peak amplitude, total energy, principal components, etc.) using MClust (offline spike sorting software, written by A. D. Redish). Only well-separated single units defined by isolation distance > 15, L-ratio < 0.2, and the spikes < 0.01 % with < 2 ms ^36^ were used in the data analysis.

## Data analysis

All data analysis was performed using custom scripts written in MATLAB (MathWorks). To assess the effects of optogenetic stimulation on the activation of mPFC neurons, neuronal activity relative to the light events was expressed in a peri-event time histogram (PETH) and a spike raster (rastergram) for each neuron. To examine light artifact in light-excited units, the similarity of spike waveform shapes for 1 sec before and after the light onset was determined by calculating the correlation coefficient (*r*) between two average spike waveforms for each unit. For spectral analysis, LFP signals were downsampled to 1 kHz. The LFP spectrogram (4-100 Hz) was computed by convolving the LFP signal with a complex Morlet wavelet. All values are given as mean ± SEM unless otherwise stated.

## Data availability

The datasets generated and/or analyzed in the current study are available from the corresponding authors upon reasonable request.

## Acknowledgements

This work was supported by a Wallenberg Academy Fellow in medicine grant (Dnr KAW 2012.0131) to M.C. from the Knut and Alice Wallenberg Foundation, the Swedish Research Council (Dnr 2016-02700), and Karolinska Institutet (Dnr 2016-00139). We thank H. Park for assistance with tissue processing.

## Author contributions

H.K. initiated this work and created the DMCdrive design. H.K. and H.S.B. designed and performed experiments. H.K., H.S.B., and M.C. analyzed data, prepared figures, and wrote the manuscript.

## Competing interests

The authors declare no competing interests.

## Notes

### Competing Interest Statement

The authors have declared no competing interest.

## References

1. Jeantet, Y. & Cho, Y. H. Design of a twin tetrode microdrive and headstage for hippocampal single unit recordings in behaving mice. J. Neurosci. Methods 129, 129–134 (2003).

2. Wilson, M. A. & McNaughton, B. L. Dynamics of the hippocampal ensemble code for space. Science (80-.). 261, 1055–1058 (1993).

3. Stosiek, C., Garaschuk, O., Holthoff, K. & Konnerth, A. In vivo two-photon calcium imaging of neuronal networks. Proc. Natl. Acad. Sci. U. S. A. 100, 7319–7324 (2003).

4. Jun, J. J. et al. Fully integrated silicon probes for high-density recording of neural activity. Nature 551, 232–236 (2017).

5. Muthuswamy, J., Okandan, M., Jain, T. & Gilletti, A. Electrostatic microactuators for precise positioning of neural microelectrodes. IEEE Trans. Biomed. Eng. 52, 1748–1755 (2005).

6. Fee, M. S. & Leonardo, A. Miniature motorized microdrive and commutator system for chronic neural recording in small animals. J. Neurosci. Methods 112, 83–94 (2001).

7. Voigts, J., Siegle, J., Pritchett, D. L. & Moore, C. I. The flexDrive: An ultra-light implant for optical control and highly parallel chronic recording of neuronal ensembles in freely moving mice. Front. Syst. Neurosci. 7, 1–9 (2013).

8. Jackson, N. et al. Long-Term Neural Recordings Using MEMS Based Movable Microelectrodes in the Brain. Front. Neuroeng. 3, 10 (2010).

9. Boyden, E. S., Zhang, F., Bamberg, E., Nagel, G. & Deisseroth, K. Millisecond-timescale, genetically targeted optical control of neural activity. Nat. Neurosci. 8, 1263–1268 (2005).

10. Kim, C. K., Adhikari, A. & Deisseroth, K. Integration of optogenetics with complementary methodologies in systems neuroscience. Nat. Rev. Neurosci. 18, 222–235 (2017).

11. Cohen, J. Y., Haesler, S., Vong, L., Lowell, B. B. & Uchida, N. Neuron-type-specific signals for reward and punishment in the ventral tegmental area. Nature 482, 85–88 (2012).

12. Lima, S. Q., Hromádka, T., Znamenskiy, P. & Zador, A. M. PINP: A new method of tagging neuronal populations for identification during in vivo electrophysiological recording. PLoS One 4, (2009).

13. Anikeeva, P. et al. Optetrode: A multichannel readout for optogenetic control in freely moving mice. Nat. Neurosci. 15, 163–170 (2012).

14. Cardin, J. A. et al. Driving fast-spiking cells induces gamma rhythm and controls sensory responses. Nature 459, 663–667 (2009).

15. Nguyen, D. P. et al. Micro-drive array for chronic in vivo recording: tetrode assembly. J. Vis. Exp. (2009). doi:10.3791/1098

16. Ung, K. & Arenkiel, B. R. Fiber-optic implantation for chronic optogenetic stimulation of brain tissue. J. Vis. Exp. e50004 (2012). doi:10.3791/50004

17. Voigts, J., Newman, J. P., Wilson, M. A. & Harnett, M. T An easy-to-assemble, robust, and lightweight drive implant for chronic tetrode recordings in freely moving animals. J. Neural Eng. (2020). doi:10.1088/1741-2552/ab77f9

18. Kim, H., Ahrlund-Richter, S., Wang, X., Deisseroth, K. & Carlen, M. Prefrontal Parvalbumin Neurons in Control of Attention. Cell 164, 208–218 (2016).

19. Kvitsiani, D. et al. Distinct behavioural and network correlates of two interneuron types in prefrontal cortex. Nature 498, 363–366 (2013).

20. Courtin, J. et al. Prefrontal parvalbumin interneurons shape neuronal activity to drive fear expression. Nature 505, 92–96 (2014).

21. Jackson, A. & Fetz, E. E. Compact movable microwire array for long-term chronic unit recording in cerebral cortex of primates. J. Neurophysiol. 98, 3109–3118 (2007).

22. Stark, E. et al. Inhibition-induced theta resonance in cortical circuits. Neuron 80, 1263–1276 (2013).

23. Buzsaki, G., Anastassiou, C. A. & Koch, C. The origin of extracellular fields and currents--EEG, ECoG, LFP and spikes. Nat. Rev. Neurosci. 13, 407–420 (2012).

24. Sohal, V. S., Zhang, F., Yizhar, O. & Deisseroth, K. Parvalbumin neurons and gamma rhythms enhance cortical circuit performance. Nature 459, 698–702 (2009).

25. Siegle, J. H., Pritchett, D. L. & Moore, C. I. Gamma-range synchronization of fastspiking interneurons can enhance detection of tactile stimuli. Nat. Neurosci. 17, 1371–1379 (2014).

26. Kim, D. et al. Distinct Roles of Parvalbumin- and Somatostatin-Expressing Interneurons in Working Memory. Neuron 92, 902–915 (2016).

27. Lee, S.-H. et al. Activation of specific interneurons improves V1 feature selectivity and visual perception. Nature 488, 379–383 (2012).

28. Pi, H.-J. et al. Cortical interneurons that specialize in disinhibitory control. Nature 503, 521–524 (2013).

29. Jackson, J., Karnani, M. M., Zemelman, B. V., Burdakov, D. & Lee, A. K. Inhibitory Control of Prefrontal Cortex by the Claustrum. Neuron 99, 1029–1039.e4 (2018).

30. Delcasso, S., Denagamage, S., Britton, Z. & Graybiel, A. M. HOPE: Hybrid-Drive Combining Optogenetics, Pharmacology and Electrophysiology. Front. Neural Circuits 12, 41 (2018).

31. Billard, M. W., Bahari, F., Kimbugwe, J., Alloway, K. D. & Gluckman, B. J. The systemDrive: a Multisite, Multiregion Microdrive with Independent Drive Axis Angling for Chronic Multimodal Systems Neuroscience Recordings in Freely Behaving Animals. eNeuro 5, (2018).

32. Yizhar, O., Fenno, L. E., Davidson, T. J., Mogri, M. & Deisseroth, K. Optogenetics in neural systems. Neuron 71, 9–34 (2011).

33. Liang, L. et al. Scalable, Lightweight, Integrated and Quick-to-Assemble (SLIQ) Hyperdrives for Functional Circuit Dissection. Front. Neural Circuits 11, 8 (2017).

34. Billard, M. W., Bahari, F., Kimbugwe, J., Alloway, K. D. & Gluckman, B. J. The systemdrive: A multisite, multiregion microdrive with independent drive axis angling for chronic multimodal systems neuroscience recordings in freely behaving animals. eNeuro 5, 1–19 (2018).

35. Calvigioni, D. et al. Functional Differentiation of Cholecystokinin-Containing Interneurons Destined for the Cerebral Cortex. Cereb. Cortex 27, 2453–2468 (2017).

36. Schmitzer-Torbert, N., Jackson, J., Henze, D., Harris, K. & Redish, A. D. Quantitative measures of cluster quality for use in extracellular recordings. Neuroscience 131, 1–11 (2005).

